# Lesions in the cerebellum impact cross-modal temporal predictions

**DOI:** 10.1101/2025.07.04.663130

**Authors:** Antonio Criscuolo, Franziska Knolle, Michael Schwartze, Erich Schröger, Richard B. Ivry, Sonja A. Kotz

## Abstract

The cerebellum (CE) supports the encoding of the precise sensory event timing and the generation of temporal predictions. Here we investigated whether focal CE lesions impact temporal predictions in a cross-modal context. Individuals with CE lesion (n=9) and healthy-matched controls (HC) were presented with visuo-auditory stimulus pairs, presented in a temporally regular (predictable) or irregular (unpredictable) manner while EEG was recorded. We hypothesized cross-modal temporal predictions to be mediated by pre-stimulus cerebello-cortical beta-band (12-25Hz) activity. In turn, we expected HC, but not CE patients, to show a modulation of pre-stimulus beta power as a function of temporal prediction.

HC showed greater pre-stimulus beta-band suppression in anticipation of sound onsets, and stronger post-stimulus delta- and theta-band (1-4Hz; 4-8Hz) power in the predictable than the unpredictable condition. Furthermore, they displayed a significant modulation of pre-stimulus delta-beta cross-frequency coupling as a function of temporal prediction. These effects were not observed in the CE group.

Results confirm that cerebellar lesions impair the generation of temporal predictions in cross-modal (visuo-auditory) stimulus processing, extending the role of cerebellar predictive timing from sensorimotor to motor-independent cross-modal perception.

## Introduction

The cerebellum (CE) is characterized by fast spiking activity (>200Hz ^1–3^) supporting subsecond temporal processes ^4–7^. Computations of cerebellar timing support the encoding of precise timing of sensory input ^8–10^ and the estimation of temporal intervals ^11–16^. In turn, these core temporal computations are exploited to generate sensorimotor and temporal predictions ^1,17–21^, e.g., in the generation and assessment of forward models ^1,18,21–23^. Hence, the CE engages in generating internal representation of the sensory consequences of motor plans, utilized in motor coordination ^1,17^ and predictive motor timing ^19–21^, and motor-to-somatosensory predictions prepare the musculoskeletal system for optimal motor control ^18,24,25^, e.g., in speech ^26,27^.

As there is strong evidence linking the CE to various non-motor high-order cognitive functions - from learning ^28^, reward processing, working memory, social cognition to language processing ^29–33^ - the question arises as to whether the CE not only generates somatosensory predictions, but modality-independent sensory predictions.

This question was previously addressed by Knolle and colleagues, who showed that patients with focal CE lesions did not adaptively suppress event-related neural responses to self-generated compared to externally-generated sounds ^34,35^. The authors concluded that the CE is involved in generating motor-to-auditory predictions, ultimately extending its role from motor-to-somatosensory to modality-independent predictions. In turn, CE lesions may ultimately impact the ability to utilize predictions to optimize performance in task-based settings ^1,36–38^. Together, these observations motivated a second question: if the CE gages temporal ^1,17–21^ and modality-independent ^34,35^ predictions, does the CE generate motor-independent temporal predictions in a cross-modal context?

In this study, we assessed *whether* focal cerebellar damage modulates sensory event encoding and temporal predictions in a cross-modal setting eliminating motor involvement. Individuals with CE lesion and healthy-matched controls (HC) were presented with paired visual (flash) and auditory (tone) stimuli, embedded in either a temporally predictable (fixed timing) or unpredictable (varied timing) context, while we recorded their EEG. If the CE gages temporal predictions in a modality independent manner, individuals with CE lesions should not not show a comparable processing advantage to HC in a temporally predictable compared to an unpredictable context.

To address this question, we targeted two neural signatures typically associated with sensory and temporal predictions: event-related beta (β; 12-25Hz) and delta (δ; 1-4Hz) activity.

In temporally regular contexts (e.g., isochronous sensory stimulation), β-activity in cerebello-cortical networks encodes the timing of sensory input and conveys temporal predictions ^21,39–42^ in anticipation of action initiation ^43^ and sensory onsets ^44^. Thus, β-power displays precise event-related temporal dynamics: β-power is suppressed (also referred to as β-desynchronization) prior to stimulus onset, and reaches a positive power peak post-stimulus (β-synchronization) ^44–46^. Pre-stimulus and movement ^47^ β-suppression is thought to reflect a predictive mechanism associated with the encoding and retention of predictable temporal structure ^48–51^. The strength of event-related cerebellar β-dynamics can be modulated by temporal predictability in sensory streams ^52^: Andersen and colleagues showed that cerebellar β-strength is higher in temporally regular than irregular contexts, supporting its role in maintaining an internal representation of the sensory environment and conveying predictions. Furthermore, β-dynamics are thought to drive top-down modulation of sensory regions ^53,54^, thus coupling with bottom-up post-stimulus δ-band activity ^55– 58^. This view is substantiated by clinical studies showing that CE lesions alter the capacities to generate and maintain stable neural representations of temporal regularities in the sensory environment ^59,60^. For instance, Criscuolo and colleagues showed that CE patients with focal CE lesions failed to encode temporally regular sound trains: differently from HC, individuals with CE lesions had difficulties in *tuning* δ-activity to sound onsets, showing a compromised ability to encode, track, and predict rhythmic sound onsets ^59^. These individuals further displayed altered δ dynamics when estimating temporal intervals ^61^, further supporting the role of the CE and δ in conveying temporal predictions.

Given the role of β-δ bands in temporal event encoding and temporal prediction, this study assessed *whether* CE lesions alter event-related neural dynamics in the β-δ bands, and β-δ cross-frequency coupling in temporally variable cross-modal contexts.

## 2. Materials & Methods

### 2.1. Participants

Eighteen participants took part in the study and signed written informed consent (IRB 410-14-15122014 in accordance with ethics committee of the University of Leipzig, Germany) and the declaration of Helsinki. One group comprised nine individuals (CE: mean age 48, range = 30-71 years; 5 males) with focal cerebellar lesions due to either a cerebrovascular event (n=8) or tumor resection (n=1). Demographic information along with etiology of neurological event are provided in Table 1, along with lesion reconstructions in Figure 1. Nine neurologically healthy controls, matched for age, gender, and education formed the control group (HC: mean age 52.1, range = 28-63 years; 5 males). The HC group was recruited via a participant database at the Max Planck Institute for Human Cognitive and Brain Sciences (Leipzig, Germany). All participants were right-handed and reported no history of hearing or psychiatric disorders. All participants received monetary compensation for taking part in the study.

**Table 1.** Information on individuals’ demographics and cerebellar lesion. The table provides, in order: Participant number, gender, age, lesion location, lesion aethiology. Acronyms: M/F = male or female; SCA = superior cerebellar artery; PICA = posterior inferior cerebellar artery; SAB = subarachnoid hemorrhage.

| **Patient N** | **Gender** | **Age** | **Lesion Site** | **Aetiology** |
| --- | --- | --- | --- | --- |
| 1 | M | 46 | Pedunculus cerebellaris superior and inferior; Lobus cerebellaris anterior | SCA infarction; SAB |
| 2 | M | 71 | Lobus posterior cerebellum | PICA infarction |
| 3 | M | 60 | Inferior lobus cerebellum posterior | PICA infarction |
| 4 | F | 39 | Posterior cerebellar lobule | PICA infarction |
| 5 | F | 46 | Bilateral inferomedial; Vermis 2 | PICA infarction |
| 6 | M | 55 | Left posterior cerebellar lobule | PICA infarction |
| 7 | F | 33 | Bilateral Vermis; superior cerebellar peduncle | tumor |
| 8 | M | 30 | Left lateral cerebellar lobule; posterior cerebellar lobule | intracerebellar hemorrhage |
| 9 | F | 50 | Right anterior cerebellar lobule; medial cerebellar lobule | PICA infarction |

**Figure 1.**
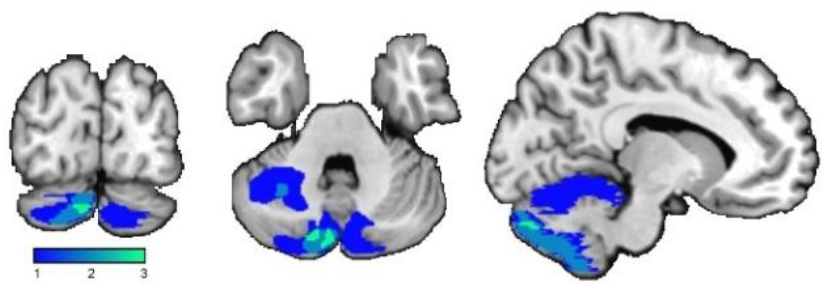
Brain lesion delineation and overlap. The figure provides the brain lesion delineation on a template anatomical MRI for individuals with cerebellar (CE) lesion. The figures were obtained by using MRIcron (http://www.mccauslandcenter.sc.edu/mricro/mricron/), and display the lesion overlap color-coded in shades of blue (blue = minimal overlap; light blue high overlap; values represent the N of individuals in a range from 0 to 3).

**Figure 2.**
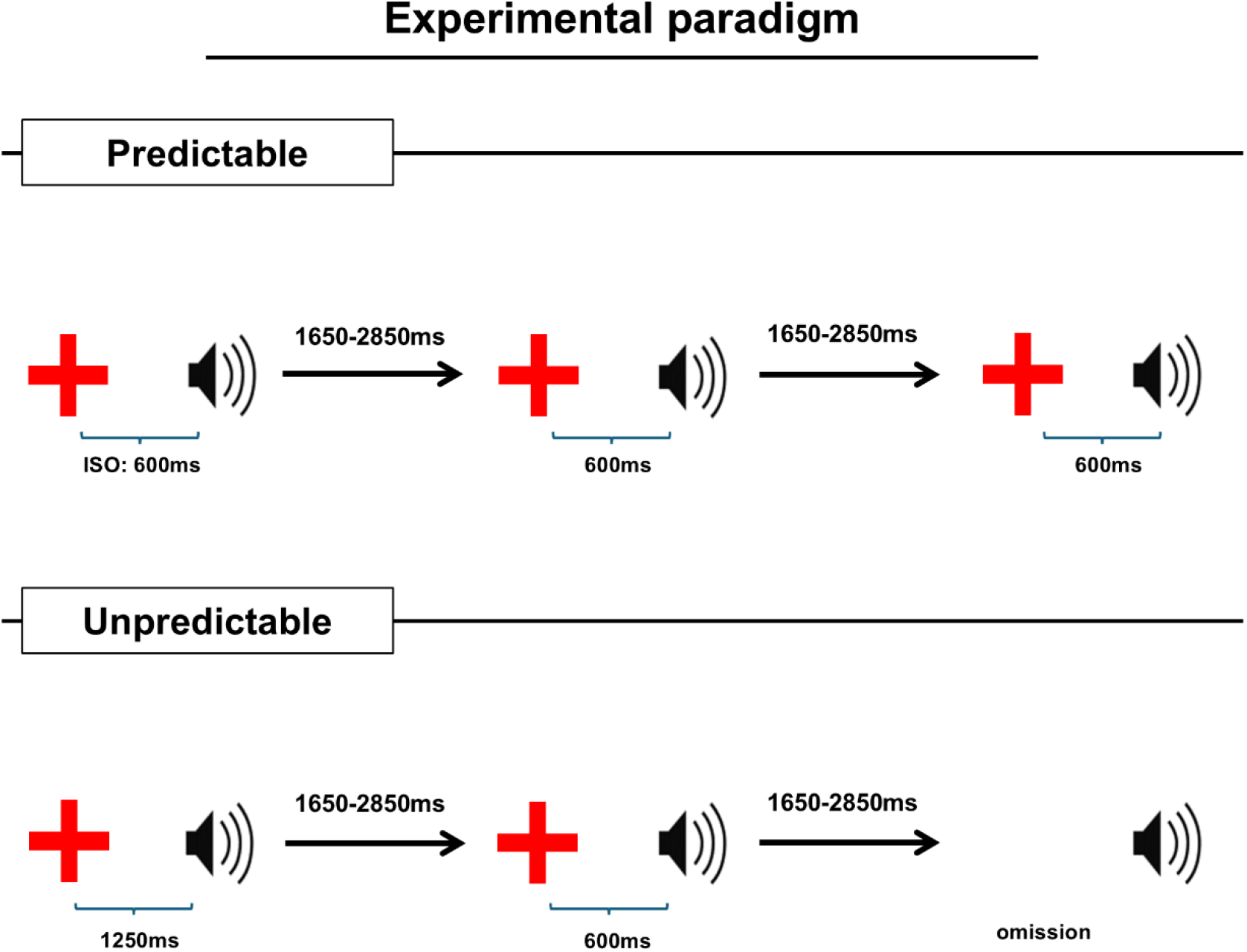
Experimental paradigm. Participants listened to stimulus pairs consisting of different color crosses (visual cue) and a sinusoidal tone. The inter-stimulus onset (ISO) was fixed at 600ms in the predictable condition (top), while varied between 450, 600, 850, 1250, or 1650ms in the unpredictable condition (bottom). The visual cue was omitted in one third of the trials in the unpredictable condition. The intertrial interval ranged between 1650-2850ms.

### 2.2. EEG experiment: design and procedure

We used a passive listening task in which participants were presented with pairs of of visuo-auditory stimuli, but did not have to perform a response. There were two conditions, each containing two blocks. All blocks were randomized. In the temporally predictable condition, a visual stimulus (a colored cross, randomly occurring in one of five colors) was presented for 150 ms. After a 600ms interstimulus interval (ISI on all predictable trials) a visual stimulus a sinusoidal tone was played (F0 = 680Hz, duration = 50ms, amplitude= 70dB SPL). Thus, the onset of the visual cue predicted the temporal onset of the tone. Participants performed two blocks of 75 trials each, with a total of 150 trials. Within each block, each colored cross appeared 15 times. The duration of the inter-trial interval (ITI) was randomly selected from a uniform distribution ranging from 1650 to 2850 ms. Thus, there was no temporal predictability from one trial to the next.

In the temporally unpredictable condition, participants performed two blocks of 225 trials each and the duration of the ITI again ranged from 1650 to 2850 ms. The 225 trials per block consisted of 75 trials with an ISO of 600ms, 75 trials with an ISO of 450 ms, 850 ms, 1250 ms, or 1650 ms, randomly selected, and 75 trials in which the visual stimulus was eliminated (tone only). In the temporally unpredictable condition, participants could not predict the onset time of the auditory stimulus. In total, the task contained 600 trials.

To evaluate EEG signatures of temporal prediction, we compared neural responses in the predictable and unpredictable conditions with the same ISO: 600ms. With a fixed ISO across conditions, the only variable being manipulated was the temporal predictability. The visuo-auditory pairings with a 600 ms ISO was presented 150 times for the both the predictable and unpredictable condition. All subsequent analyses focussed on these trials. Of note, participants did not perform a task, and did not respond to any stimulus onset. The absence of a task was chosen to eliminate any EEG responses that could be related to motor readiness/anticipation of an action.

#### 2.2.1. EEG recording

EEG was recorded from 55 Ag/AgCl scalp electrodes positioned according to the International 10-10 system with the ground placed on the sternum. Four additional vertical and horizontal electrodes monitored eye movements and were positioned on the outer canthus of each eye, and on the inferior and superior areas of the left orbit. The signals were amplified, digitized using a sampling rate of 500Hz (Refa amplifiers system, TMS international, Enschede, NL) and an anti-aliasing filter of 135 Hz was applied. Electrode impedances were kept below 5kΩ and the left mastoid served as online reference. Data was referenced to the left and right mastoids offline.

#### 2.2.2. Data Analysis

##### EEG Preprocessing

EEG data were analyzed in MATLAB combining custom scripts and the FieldTrip toolbox 62. Data were band-pass filtered with a 4th order Butterworth filter in the frequency range of 0.1-50Hz (*ft_preprocessing*). Eye-blinks and other artifacts were identified using independent component analysis. Components with a strong correlation (>.4) with the EOG time-courses were automatically identified and removed with ‘*ft_rejectcomponent*’ before reconstructing the EEG time-course. Components were visually inspected to ensure removal of blinks and cardiac artifacts. On average, 2 components were removed (‘*ft_rejectcomponent*’) before reconstructing the EEG time-series. We employed artifact subspace reconstruction (‘*pop_clean_rawdata*’ function in EEGlab) and an artifact suppression procedure. The artifact suppression procedure ^50,51,59,63^ interpolated noisy (>100uV) time-windows on a channel-by-channel basis. Data were low-pass filtered at 40Hz via ‘*ft_preprocessing*’, segmented relative to each tone onset (-2 to +2s), and downsampled to 250Hz. As in prior work ^50,51,59,63^, subsequent analyses focused on a fronto-central channel (FC) cluster, encompassing the sensor-level correspondents of prefrontal, pre-, para-, and post-central regions engaged in auditory and temporal processing ^64,65^. The cluster included 14 channels: ‘AFz’, ‘AF3’, ‘AF4’, ‘F3’, ‘F4’, ‘F5’, ‘F6’, ‘FCz’, ‘FC3’, ‘FC4’, ‘FC5’, ‘FC6’, ‘C3’, ‘C4’.

##### Time-frequency analyses

After preprocessing, we performed time-frequency transformation (‘*ft_freqanalysis’*) by means of a wavelet-transform ^66^. The bandwidth of interest was set between 1-30Hz, using a frequency resolution of 1Hz. The number of fitted cycles was set to 3 for lower frequencies (up to 4Hz) and to 7 for higher frequencies (above 4Hz). No averaging over channels, trials, or participants was performed at this stage. Data were then re-segmented from -800 to 600ms relative to tone onset and standardized using (z-scored normalization).

We statistically assessed the effect of temporal prediction within each group by comparing EEG signatures of interest in the predictable and unpredictable conditions. Statistical significance was tested via Monte Carlo cluster-based permutation testing (‘ft_freqstatistics’). We focused on a time-window extending from -400 to 400ms relative to tone onset, thus avoiding visual evoked neural activity (-600ms). We used an alpha level of .025, 500 randomizations, and a minimum cluster size of 2 neighboring channels.

We assessed group differences by comparing group differences by comparing EEG signatures of temporal prediction in the CE and HC groups. First, we used ‘ft_math’ to calculate the difference between neural responses to temporally predictable and unpredictable conditions at the individual level. Next, we calculated the group average by using ‘ft_grandaverage’. Finally, we compared group differences via Monte Carlo permutation testing (‘ft_freqstatistics’). We focused on a time-window extending from -400 to 400ms relative to tone onset, using an alpha level of .025, 500 randomizations, and a minimum cluster size of 2 neighboring channels. Because both within-subject and between-group analyses utilized a 2-sided test, the alpha level was adjusted to distribute the probability over both tails.

##### Cross-frequency coupling

To perform cross-frequency coupling (CFC) analyses, we performed time-frequency analyses as described above, but this time extracted complex exponentials to calculate delta (1-4Hz) - beta (12-25Hz) CFC. To this end, we employed the ‘ft_crossfrequencyanalysis’ function in FieldTrip. Coherence is calculated as:

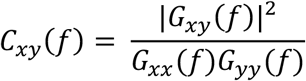

where the numerator features the cross-spectral density between x and y, and the denominator the auto spectral density of x and y, respectively. Coherence is a value between 0 and 1, defining how much we can predict time series y based upon time series x.

To obtain a dynamic measure of coherence (as compared to a single summary value across the entire time-segment of interest), we used a 100ms sliding window over the range from -600 to 600ms relative to sound onset in steps of 50ms.

We assessed group differences via Monte Carlo cluster-based permutation testing. We focused on a time-window extending from -400 to 400ms relative to tone onset, using an alpha level of .025 and 500 randomizations.

### 2.3. Data and code Availability

The analysis code in use here will be provided as open access resource after publication. Lesion data and participant data in use are under data protection rules.

## 3. Results

### 3.1. Time-frequency analyses

We used time-frequency analyses to explore (i) neural signatures of cross-modal sensory temporal prediction and (ii) how these signatures are impacted in inviduals with focal cerebellar lesions.

HC showed a stronger pre-stimulus beta suppression (-.4 to 0s; 18-24Hz) in the temporally predictable as compared to unpreditable conditions (Fig. 3A, right), while individuals with CE lesions did not show this effect. Further, HC displayed a stronger post-stimulus delta-(1-4Hz) and theta (4-8Hz) band power increase (0 to .4s) in the temporally predictable than the unpredictable condition (also see Suppl. Fig. 1). This latter effect seemed reversed in individuals with CE lesions, but within-subject analyses per group showed that the temporal prediction effect did not yield a significant power modulation in the CE group (Suppl. Fig. 1).

**Figure 3.**
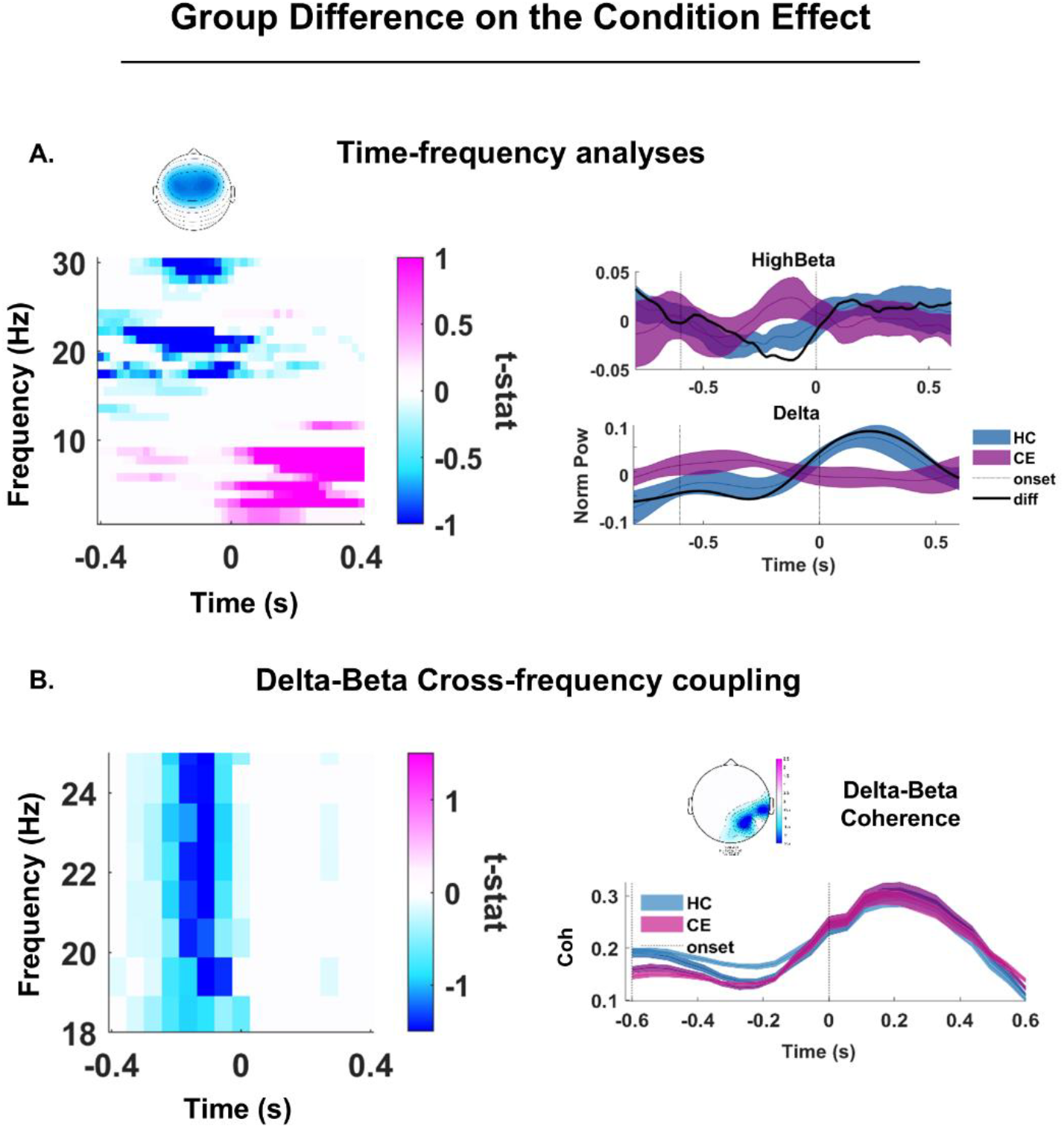
Time-frequency analyses on the predictability effect. A: Group difference (HC – CE) for the temporal prediction effect (Predictable – Unpredictable). The plot features time on the x-axis (-.4 to .4s relative to sound onset) and frequency (1-30Hz) on the y-axis. Blue for negative and pink for positive t-values. The plot is masked by the probability map obtained via Montecarlo permutation testing: in color, all significant results (p < .025). At the top, the delineation of the fronto-central channel cluster of interest. On the right, the time-course of the temporal prediction effect (Predictable – Unpredictable), in blue for HC and purple for CE. Solid line reports the mean, while the shades display the standard error across participants. The dark link is the group difference (HC – CE). Top plot: high-beta activity (20-25Hz); bottom: delta activity (1-4Hz). The plot features time on the x-axis (-.8 to .6s relative to sound onset) and normalized power (z-value) difference on the y-axis. Vertical dotted lines indicate the visual cue (-.6s) and sound (0s) onset, respectively. B: group difference (HC – CE) for the temporal prediction effect (Predictable – Unpredictable) in delta-beta cross frequency coupling. The plot features time on the x-axis (-.4 to .4s relative to sound onset) and frequency (18-30Hz) on the y-axis. Blue for negative and pink for positive t-values. The plot is masked by the probability map obtained via Montecarlo permutation testing: in color, all significant results (p < .025). On the right, the time-course of the temporal prediction effect (Predictable – Unpredictable), in blue for HC and purple for CE (dark color for Predictable and light color for Unpredictable). Solid line reports the mean, while the shades display the standard error across participants. The plot features time on the x-axis (-.6 to .6s relative to sound onset) and coherence difference on the y-axis. Vertical dotted lines indicate the visual cue (-.6s) and sound (0s) onset, respectively. The topographic plot on top displays the sensor-level map of significant effects in the pre-stimulus interval (-.3s to 0).

### 3.2. Cross-frequency coupling

Next to power changes, we hypothesized that predictive neural responses in anticipation of a tone onset should also display a modulation in delta-beta cross-frequency coupling (CFC).

We observed one predominant effect: HC showed a more enhanced pre-stimulus CFC suppression in the temporally predictable (darker blue) than the unpredictable (lighter blue) condition (Fig. 3B, right). The pre-stimulus CFC effect was predominant over a right electrode cluster (Fig. 3B, top right). The effect of temporal prediction was absent in individuals with CE lesions (Fig. 3B).

## 4. Discussion

In this study, we aimed to investigate whether focal cerebellar modulate temporal predictions in a cross-modal, task-free (non-motor) setting. Individuals with CE lesions and healthy-matched controls (HC) were exposed to visuo-auditory stimulus pairs presented in a temporally predictable or unpredictable condition.

We expected HC, but less so individuals with CE lesions, to efficiently encode temporal events and generate temporal predictions in anticipation of a sound onset in these visuo-auditory stimulus pairs. Thus, we expected to observe a modulation of event-locked β-δ neural responses as a function of temporal prediction in HC, but expected this to be changed or even absent in individuals with CE lesions.

Time-frequency analyses confirmed group differences for the temporal prediction effect: HC, but not individuals with CE lesions, showed a modulation of pre-stimulus high-β (18-25Hz) and post-stimulus δ- and theta-band (1-4Hz; θ, 4-8Hz) power as a function of temporal prediction. More specifically, in HC β-power showed a stronger pre-stimulus suppression in anticipation of a sound onset in the temporally predictable than the unpredictable condition. Similarly, HC showed stronger δ- and θ-band responses in the temporally predictable than the unpredictable condition post-stimulus. Cross-frequency coupling (CFC) analyses further revealed a significant modulation of β-δ coherence prior to stimulus onset as a function of temporal prediction in the HC. The modulatory effect of temporal prediction on event-locked β-δ and CFC was nearly absent in the CE lesion group.

These results further confirm the fundamental role of the CE in forming internal representations of the temporal structure of the sensory environment ^1,18,21–23^, enabling estimates and predictions of temporal intervals ^11–16^. The current data also support the notion that CE lesions minimize the use of temporal cues to enhance senstitivity to specific moments in time ^67^, as indexed by reduced δ-dynamics ^59,61^. Similarly, the absence of a post-stimulus θ-modulation as a function of temporal prediction might support the role of θ in the temporal sampling of sensory input ^68^, and the generation of a temporally structured representation of sensory signals that facilitates perception ^69^.

Our findings further substantiate the role of cerebellar β-activity in mediating temporal predictions ^21,39–42^, and adaptively modulating β-strength as a function of temporal predictability ^52^. Finally, the anticipatory β-δ CFC modulation observed in HC is indicative of β’s role in driving top-down modulation of sensory regions ^53,54^ by coupling with bottom-up δ ^55–58^.

While the CFC results were observed across the entire β-band, in HC the pre-stimulus modulation (temporal prediction) was most evident in the upper β-band (18-25Hz). Although typically treated as a homogenous frequency-band, low- and high-β frequencies can be differentiated into motor preparation and temporal prediction, respectively ^70,71^. Finding pre-stimulus upper-β band acitivty might support such a dissociation and strengthen the link between pre-stimulus high-β and temporal prediction ^44,72–75^, also in a cross-modal context. Together, these results not only strengthen the notion that CE lesions can impair the ability to exploit temporal predictions to optimize sensory processing ^1,36–38^, but fundamentally extend the role of the CE in generating non-motor temporal predictions in a cross-modal context.

Given the role of the CE in sensorimotor and temporal predictions ^1,17–21^, as well as in modality-independent ^34,35^ predictions, can one consider whether the CE is involved in domain-general temporal predictions? This question is at the core of an ongoing debate: does the CE perform domain-general universal computations ^3^, or does it rely on domain- and task-specific functional ones ^76^?

Within the extended cerebello-cortico and cortico-basal ganglia-cerebellar networks ^3,10,21,23,28,77–83^, the CE does not only perform temporal computations ^8,10,21,84^, but is coordinates motor and sensorimotor predictions ^1,17–21^, learning ^28^ and general cognition, spanning from reward processing, working memory, social cognition to language processing ^29–33^. Furthermore, structural and functional change in the CE has been linked to various neurodevelopmental ^85^, neurological ^86^, and psychiatric ^1,87–89^ disorders, ranging from autism ^90,91^, epilepsy ^92^, to psychosis ^87,93,94^. Altogether, basic and translational research suggest the CE to play a fundamental role in neurocognitive functions, from health to pathology. Yet, it remains to be better elucidates *why* this is the case: altered timing and predictions seem to be good candidates to explain the pathophysiology of some psychotic and psychiatric symptoms ^89,95^; but more research is needed to better characterize CE neuro-anatomical, -functional, and physiological properties and their link to perception, action, and cognition. Thus, we encourage future research to consider the contribution of the CE in monitoring cortical functions ^96^.

While we show novel results on the role of the CE in gaging temporal predictions in a cross-modal context, we also acknowledge that this study has several limitations: first, we can only describe scalp-level activity (as opposed to source-level activity) recorded by a limited number of electrodes, which only allows for tentative conclusions on a more general functional specification of the CE within cortico-subcortical networks. Second, the absence of a behavioral task prevents drawing any direct conclusions between the observed neural activity and actual performance: e.g., we cannot ensure whether the lack of pre-stimulus beta-band modulation in the CE group would translate to a poorer perceptual performance (e.g., reaction times). However, the involvement of a task such as a button press on catch trials would have induced motor activity, which, in turn, would have confounded the results. Finally, a limited sample size and diverse lesions sites in the CE group limit the generalizability of the current results.

Within such limits, the current study nevertheless offers novel insight into the role of the CE in temporal and content predictions and associated event-related neural dynamics.

## 5. Conclusions

With a striking count of >60 billion neurons, the cerebellum (CE) contains ∼80% of neural cells in the brain. While known as the *little brain* (from Latin), we have shown that the CE gages the generation and exploitation of temporal predictions to optimize cross-modal processing. In fact, EEG activity seems to suggest that individuals with focal CE lesions fail to utilize precise temporal event coding and temporal prediction in cross-modal processing. Our results confirm and expand previous evidence on the role of the CE in basic temporal computations.

## Suppl. Figure

**Suppl. Figure 1.**
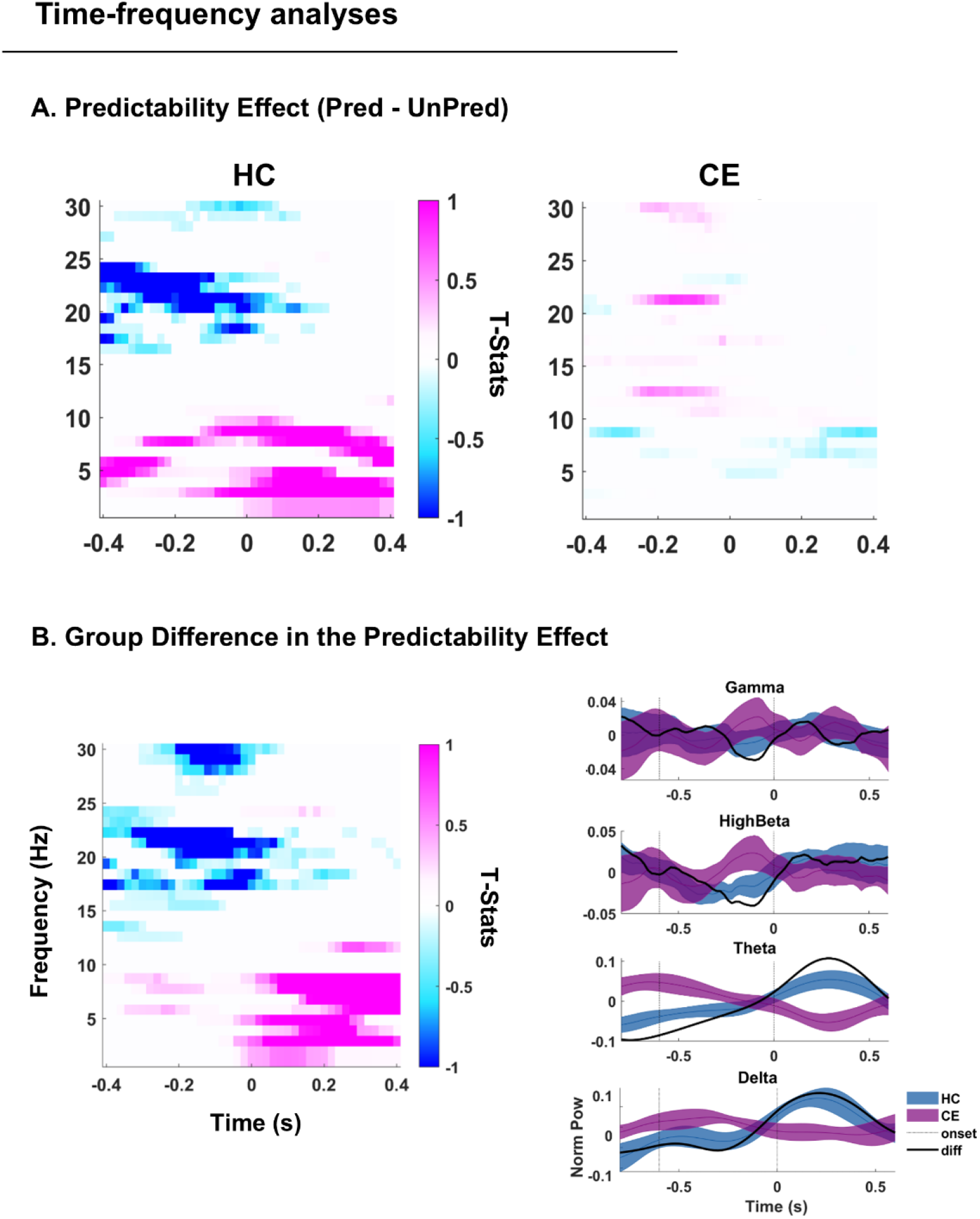
Predictability effect: within-group and between-group analyses. A: within-group analyses on the predictability (Predictable – Unpredictable) effect. The plot features time on the x-axis (-.4 to .4s relative to sound onset) and frequency (1-30Hz) on the y-axis. Left plot for HC and right plot for CE patients. Blue for negative and pink for positive t-values. The plots are masked by the probability maps obtained via Monte Carlo permutation testing: in color, all significant results (p < .025) indicated as t-values. B: group difference (HC – CE) in the predictability effect (Predictable – Unpredictable). Left plot as in A. On the right, the time-course of the predictability effect (Predictable – Unpredictable), in blue for HC and purple for CE patients. Solid line reports the mean, while the shades display the standard error across participants. The dark link is the group difference (HC – CE). Top to bottom plots: gamma (25-30Hz,) high-beta (20-25Hz), theta (4-8Hz) and delta (1-4Hz) activity. The plot features time on the x-axis (-.8 to .6s relative to sound onset) and normalized power (z-value) difference on the y-axis. Vertical dotted lines indicate the visual cue (-.6s) and sound (0s) onset, respectively.

